# Omicron Spike protein has a positive electrostatic surface that promotes ACE2 recognition and antibody escape

**DOI:** 10.1101/2022.02.13.480261

**Authors:** Hin Hark Gan, John Zinno, Fabio Piano, Kristin C. Gunsalus

## Abstract

High transmissibility is a hallmark of the Omicron variant of SARS-CoV-2. Understanding the molecular determinants of Omicron’s transmissibility will impact development of intervention strategies. Here we map the electrostatic potential surface of the Spike protein to show that major SARS-CoV-2 variants have accumulated positive charges in solvent-exposed regions of the Spike protein, especially its ACE2-binding interface. Significantly, the Omicron Spike-ACE2 complex has complementary electrostatic surfaces. In contrast, interfaces between Omicron and neutralizing antibodies tend to have similar positively charged surfaces. Structural modeling demonstrates that the electrostatic property of Omicron’s Spike receptor binding domain (S RBD) plays a role in enhancing ACE2 recognition and destabilizing Spike-antibody complexes. Collectively, our structural analysis implies that Omicron S RBD interaction interfaces have been optimized to simultaneously promote access to human ACE2 receptors and evade antibodies. These findings suggest that electrostatic interactions are a major contributing factor for increased Omicron transmissibility relative to other variants.

## Introduction

The emergence of highly transmissible Omicron variant of SARS-CoV-2 challenges our understanding of the underlying causes of its spread and clinical manifestations. Omicron is characterized by a jump in the number mutations in the crucial Spike protein with thirty amino acid changes, three small deletions, and one small insertion. In contrast, the Delta variant only has eight amino acid changes, a single amino acid deletion, and a small deletion in Spike. Indeed, tracking of SAR-CoV-2 mutations (nextstrain.org database^1^) indicates that the number of mutations in variants has been increasing over time. Since viral transmissibility also has been increasing throughout the COVID-19 pandemic, it is intriguing to speculate whether there is a mutational pattern that can provide some insight into the trajectory of emerging variants. Deciphering the links between mutations, variant evolution and transmissibility could help anticipate new variants and assist design of effective therapeutics.

Mutations in the Spike protein of the Omicron variant (BA.1 lineage) are in three major regions: two in the Spike receptor binding domain (S RBD, residues 319–541) and one in the “fusion” domain^2^. Crucially, a cluster of ten mutations is in the Spike receptor binding motif (S RBM, residues 440–505) or its ACE2-binding interface. There is also a smaller cluster of four mutations around the S1/S2 (or furin) cleavage site. The Spike N-terminal domain (S NTD, residues 13-303) contains three deletions and an insertion in addition to four point mutations. The two other Omicron lineages (BA.2 and BA.3) share twenty point mutations in Spike with the BA.1 lineage, eight of which are in S RBM. Unlike the previous SARS-CoV-2 variants, mutations in Omicron lineages tend to occur in clusters, a scenario suggested in a structural modeling analysis^3^.

The occurrence of mutation clusters challenges our understanding about their influence on the biological properties of Omicron such as antibody resistance^4-6^, mode of host cell entry^7^, viral evolution^2^, virulence^8^, and transmission. Several large-scale experimental studies have demonstrated that Omicron S RBD can evade most neutralizing antibodies, especially those targeting S RBM including some therapeutic monoclonal antibodies^4,5^. Although some Omicron S RBD mutations found in other variants are known to evade antibodies (K417N, N440K, E484K/Q), the roles of Omicron-specific mutations are not clear. According to a recent study^2^, some novel Omicron mutations (G339D, S371L, S373P, S375F, Y505H) have a negligible effect on the strength of antibody escape; such mutations are also considered rare in sarbecoviruses (SARS-CoV-1/2) and not positively selected. Antibody escape potential of multiple, clustered mutations cannot be easily assessed based on single mutation data because of likely significant conformational changes at antibody binding sites. These considerations suggest a need for new approaches to analysis of Omicron mutations, both individually and collectively.

A feature of Omicron and other variants is that most of their defining mutations are found on the surface of the Spike protein. Surface mutations affect Spike’s interactions with antibodies, ACE2 receptors and proteases to determine infectivity. To decipher the global effects of Omicron mutations, we map the electrostatic potential surface of Spike. We show that the Omicron Spike trimer surface has transitioned to a strongly positive electrostatic surface relative to the reference (Wuhan-Hu-1) Spike, especially in S RBM, which interfaces with the negatively charged ACE2 receptor. In addition, we use structural modeling to demonstrate that Omicron S RBD is attracted to ACE2 receptors by long-range electrostatic forces and can destabilize five out of six representative neutralizing antibodies tested. Thus, our computational analyses suggest that Omicron mutations collectively enhance ACE2 recognition and antibody escape.

## Results

### Spike structure of Omicron has a predominantly positive electrostatic potential surface

Use of electrostatic potential surfaces is a common approach for mapping complementary interaction interfaces in biomolecular complexes^9,10^. Relative to the reference Spike, the electrostatic surface of the Omicron Spike protein shows a marked transition to positive surface charges, especially on the top face or S RBM and near the furin (S1/S2) cleavage site (Fig. 1a); the stem region too has become less negatively charged. The additional positive surface charges in Omicron S RBM are acquired through N440K, T478K, Q493R, Q498R and Y505H mutations; the solvent-accessible loop containing the furin cleavage site added two positive charges from N679K and P681H mutations.

**Fig. 1.**
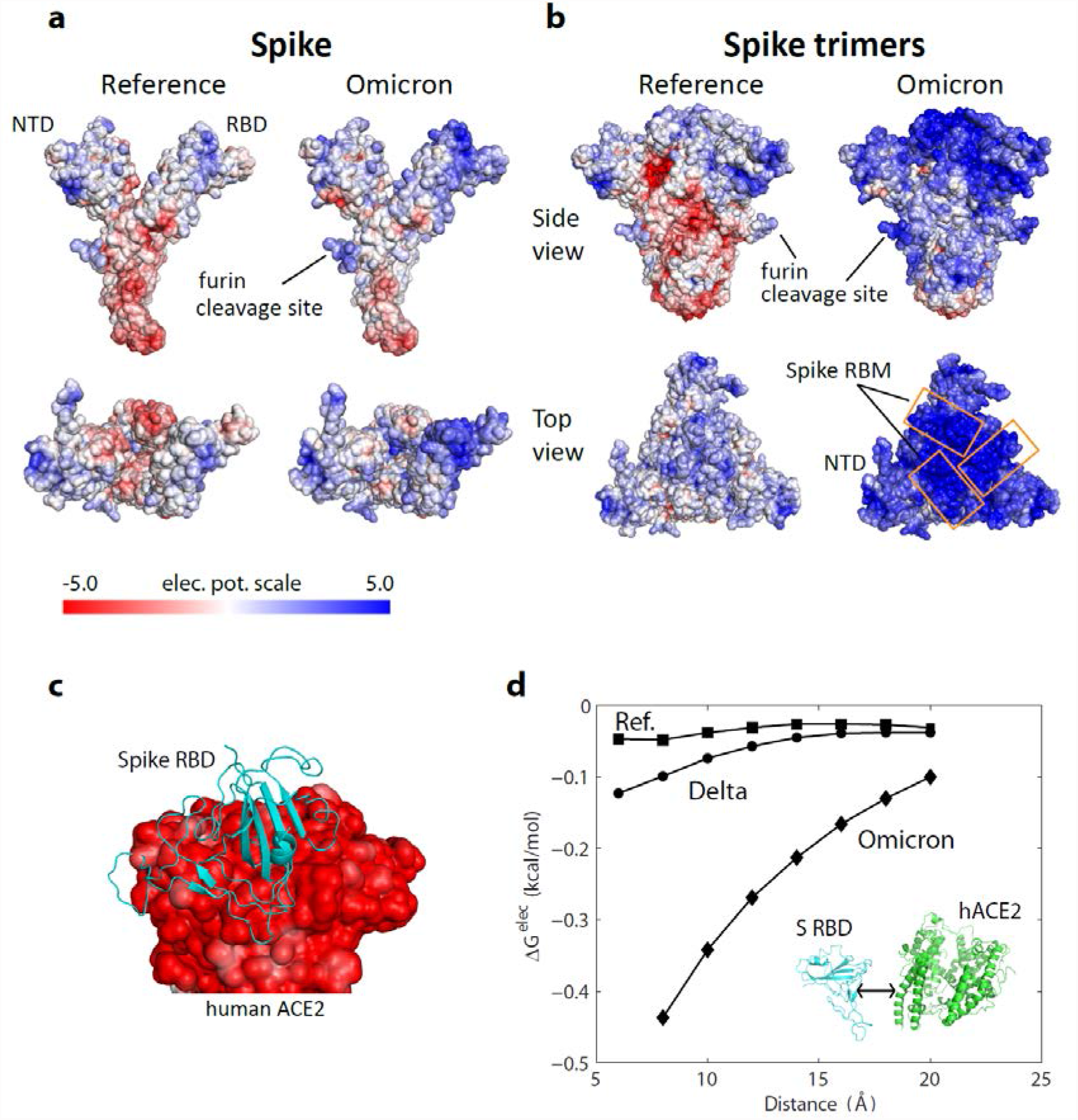
Electrostatic potential surfaces of Spike, Spike trimer, and ACE2 proteins. **a** Electrostatic surfaces of reference and modeled Omicron Spike proteins. **b** Electrostatic surfaces of reference and modeled Omicron Spike trimers. **c** Electrostatic surface of extracellular domain of human ACE2 with Spike RBD (PDB ID: 6m17). **d** Electrostatic binding energy as a function of separation distance for the reference, Delta and Omicron Spike RBD-ACE2 complexes. Separation distance is along the axis perpendicular to the interaction interface.

Since the Spike protein is assembled into Spike trimers on the viral membrane of SARS-CoV-2, the electrostatic surface of Spike trimer is a direct indication of its functional implications. The reference Spike trimer complex has a slightly positive head or top region, consisting of S NTD and RBD, while the surface of the stem region is mostly negatively charged (Fig. 1b, side view). In sharp contrast, the Omicron Spike trimer has a strongly positive head and stem regions. The Omicron Spike trimer was obtained using a predicted Spike structure (Methods) and then assembled using a solved reference Spike trimer template (PDB ID: 6vyb); a recent cryo-EM trimer structure shows a similar tertiary organization to those from reference and other variants^11^ (modeled Spike differ from the solved structure by 2.2Å and S RBD by 1.1 Å). The change in the electrostatic surface of Omicron Spike trimer is especially dramatic at the ACE2-binding interface or S RBM (Fig. 1b, top view). Since the extracellular domain of human ACE2 (residues 1-614) has an entirely negative electrostatic potential surface (Fig. 1c), Omicron S RBD is strongly attracted to target ACE2 receptors by long-range electrostatic forces.

We quantified the electrostatic interactions between S RBD and the extracellular domain of human ACE2 by computing their electrostatic binding energy as a function of separation distance for reference, Delta and Omicron Spike proteins (Fig. 1d). To simulate physiological conditions, the energies were computed using APBS v1.5 at 150 mM monovalent ions^12^. The computed electrostatic binding energies show that Omicron S RBD has a considerably greater affinity for ACE2 over all distances (8-20 Å) compared with Delta and reference S RBDs (Fig. 1d). Quantitatively, over the separation distances compared, Omicron’s electrostatic energies are 3-9 times greater than those for the reference S RBD and 3-5 times greater than Delta S RBD. Even at a distance of 20 Å Omicron S RBD is still influenced by the attractive electrostatic force emanating from the negatively charged extracellular ACE2 domain. The attractive force is expected to be even stronger if the Omicron Spike trimer, instead of S RBD, is considered because of the combined positive charges on the Spike trimer’s surface. These findings suggest that Omicron Spike trimers are guided by long range electrostatic forces to the vicinity of ACE2 leading to efficient recognition of the receptor.

### Chronology of SARS-CoV-2 variants shows that Spike RBM has accumulated positive surface charges

The advantage conferred by Omicron’s electrostatic property to infect host cells may indicate an adaptive feature of variant evolution. To reveal this plausible driver of viral dynamics, we analyzed S RBM mutations, charges, and electrostatic surfaces across SARS-CoV-2 phylogeny. To reveal biological relationships, we superposed information about the electrostatic surfaces of S RBD on the phylogeny of SARS-CoV-2 variants (nextstrain.org) as a function of viral emergence (earliest sampling date, cov-lineages.org). The Nextstrain SARS-CoV-2 phylogenetic tree represents mutation lineages and assigns each clade based on its combination of signature mutations.

The evolution of the electrostatic surface shows that there is a gradual accumulation of positive surface charges in S RBD over time and that related clades have similar electrostatic surfaces (Fig. 2). To quantify the changes, we divided the surface charge transitions into two time periods: from beginning to Fall of 2020 and late 2020 to late 2021. In the first period, Alpha, Beta, Epsilon and Lambda variants acquired moderately more positive surface charges in S RBD compared with the reference Spike. More significant accumulation of positive surface charges occurred in the second period with the emergence of Gamma, Delta, Eta, Iota, Kappa and Mu variants. The large change in surface charges in the Omicron variant appears to mark another surface charge transition from those in the second time period; Omicron sub-variants BA.2 and BA.3 have essentially the same S RBD mutations associated with charge changes.

**Fig. 2.**
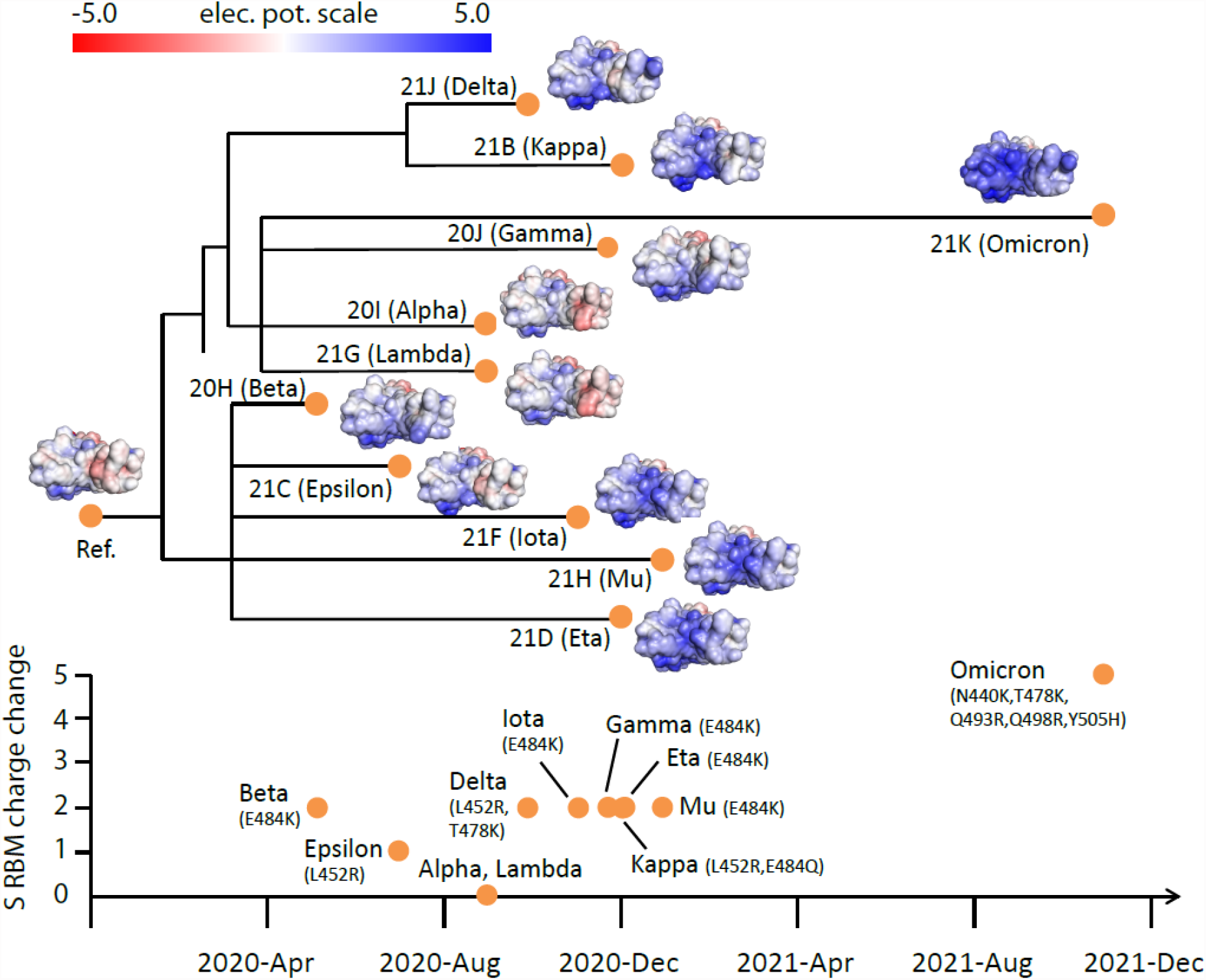
Electrostatic surfaces of Spike RBD variants organized by Nextstrain SARS-CoV-2 phylogeny. Tracking of total charge changes with accompanying mutations in S RBM suggests a gradual increase of positive charges over time.

These observations are supported by the accumulated charges in S RBM, an ACE-binding interface and target of many neutralizing antibodies. The early variants (Alpha, Beta, Epsilon and Lambda) gained an average of one elementary electric charge (+1e). In contrast, variants in the second time period (Gamma, Delta, Eta, Iota, Kappa and Mu) all gained +2e. Intriguingly, the gain in positive charges up to late 2021 is largely accounted for by only three mutations (L452R, T478K and E484K), suggesting close links between these variants. The Omicron variant gained +5e from N440K, T478K and three novel mutations Q493R, Q498R and Y505H. Thus, there is roughly a doubling of positive charges in S RBM in each of the time periods described (0→+1e→+2e→+5e).

Even though the SARS-CoV-2 clades are defined without S RBM charge considerations, their phylogeny reveals a close relationship with S RBD electrostatic surfaces. For example, Eta (clade 21D), Iota (21F) and Mu (21H) variants have similar S RBD electrostatic surfaces. Also evident is the similarly of the electrostatic surfaces of related Delta (21J) and Kappa (21B) variants. Moreover, distantly related clades (Delta/Kappa, Eta/Iota/Mu and Omicron) all acquired positive charges. These observations suggest that the increase of total positive charges in S RBM is a general adaptive feature of variant evolution to enhance ACE2 association.

### Most Omicron Spike-antibody complexes have unfavorable electrostatic interaction surfaces

Although hundreds of antibodies against SARS-CoV-2 have been sequenced, most known neutralizing antibodies target S RBD^13-15^. To assess the response of neutralizing antibodies to Omicron SARS-CoV-2, we selected six representative antibodies from high-resolution (<3 Å) S RBD-antibody complexes that bind to different S RBD sites (Fig. 3a, Methods). Specifically, antibodies P2B-2F6 and COVOX-88, 150, 158, 316 bind S RBM sites, whereas S2X259 targets outside of S RBM. Electrostatic potential surfaces of these six antibodies indicate that their Spike-interacting interfaces are predominantly positively charged, especially for P2B-2F6 and COVOX-316 (Fig. 3b). In contrast, antibody S2X259 has a negatively charged interaction interface. As shown (Fig. 3c), the antibody-binding interfaces of Spike are all positively charged. Qualitatively, this indicates that antibodies P2B-2F6 and COVOX-88, 150, 158, 316 are likely to have unfavorable electrostatic binding energies with Omicron S RBD, whereas S2X259 is expected to have a favorable binding energy. Overall, the electrostatic property of Omicron S RBD is likely to increase resistance against most classes of antibodies, as found in recent experimental studies^4,5^. Below, we present a quantitative assessment of antibody response to S RBD mutations from different variants using an antibody escape score based on binding affinity changes relative to the reference S RBD.

**Fig. 3.**
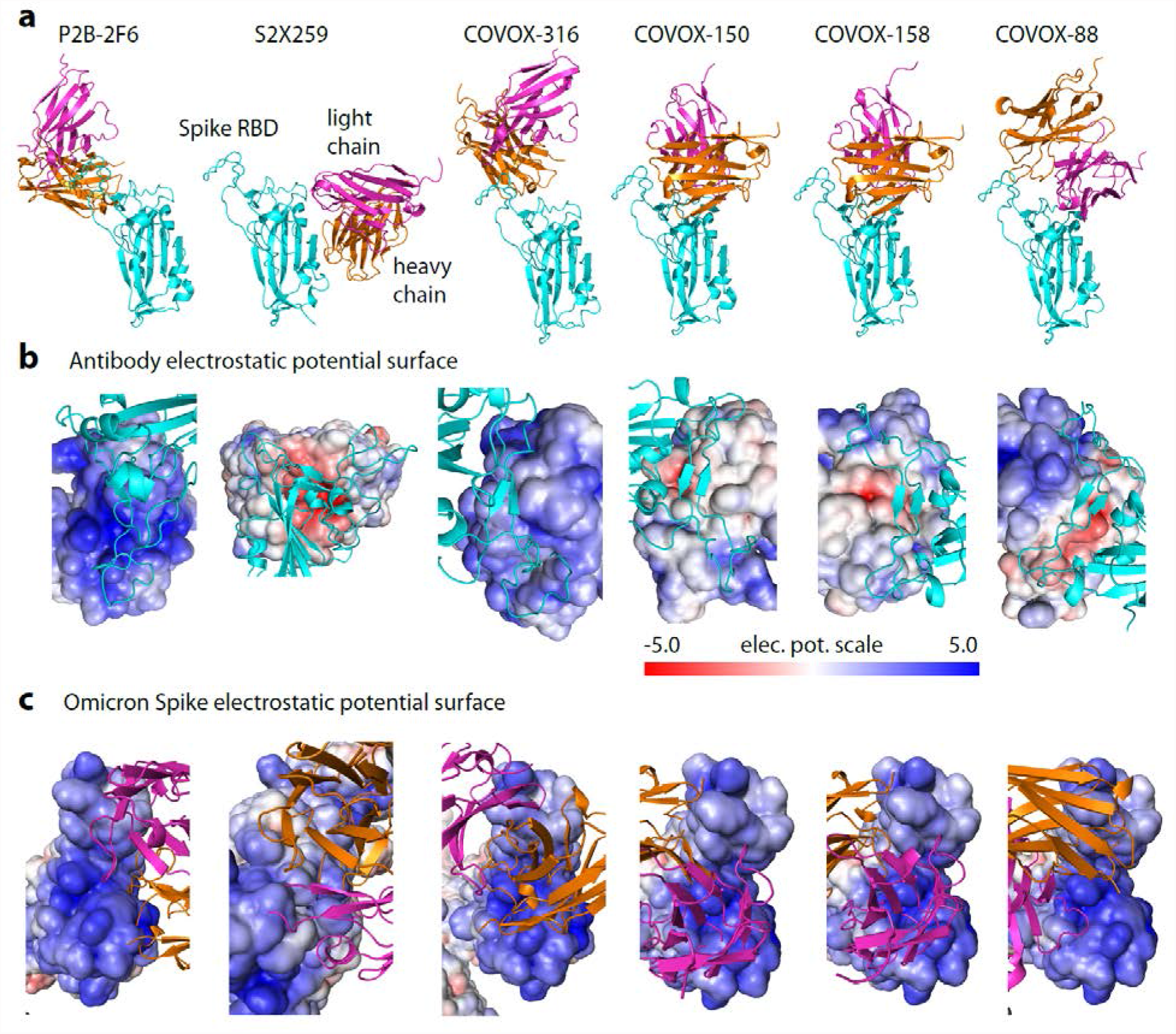
Spike-antibody complexes and their electrostatic potential surfaces. **a** Solved structures of neutralizing antibodies targeting different sites on S RBD used for computational modeling. **b** Electrostatic surfaces of antibodies complexed with reference Spike. **c** Electrostatic surfaces of Omicron Spike modeled with different antibodies using templates in (**a)**. Spike-interacting interfaces on antibodies are predominantly positively charged which contribute to antibody resistance by Omicron.

### Structural modeling captures antibody resistance of single mutations from non-Omicron variants

Current experimental approaches employ infectivity^5,16^ and yeast-display/FACS^4,17^ assays as measures of destabilization of antibody binding due to Spike mutations. Here, we use computational modeling to directly quantify the effects of Spike mutations on antibody binding. This approach is feasible because many Spike-antibody complexes have been solved in the last two years^14,15,18^, enabling structural modeling of mutational effects^19,20^. Briefly, we employed six representative neutralizing antibodies (Fig. 3, Methods) and considered mutations in direct contact with antibodies. We define the strength of antibody escape as the destabilization of antibody binding. Specifically, our antibody escape score is defined as the normalized affinity change, (ΔG_mut_ -ΔG_ref_)/|ΔG_ref_|, where ΔG_ref_ and ΔG_mut_ are affinities of reference and mutant Spike, respectively. Thus, positive scores indicate reduced antibody binding, whereas negative scores imply increased antibody binding. Here, we use the score to evaluate antibody escape induced by single mutations of the reference Spike, a method we described previously^3^.

Computed antibody escape scores show that the most prominent escape peaks across all antibodies are from E484K/Q mutations, which are confirmed escape mutants found in Beta, Gamma, Zeta and Kappa variants (Fig. 4)^17^. Mutation N501Y, occurring in Alpha, Beta and Gamma variants, only exhibits a moderate escape from COVOX-158 and COVOX-88. Mutations K417N/T from Beta and Gamma variants exhibit varying degrees of resistance to COVOX-150, COVOX-158 and COVOX-88. By contrast, S477N, a frequent mutation in GISAID database, has a weak escape from most antibodies except possibly COVOX-316. A deep mutagenesis study of single S RBD mutations based on ten antibodies determined that S477N is not an escape mutant^17^. The Delta variant has a L452Q/T487K mutation combination in S RBD. Based on our predictions, mutation L452Q can escape all tested antibodies except S2X259, and mutation T478K escapes COVOX-316 and COVOX-88. The predicted extensive antibody escape profile for the Delta variant correlates well with experimental studies^21^ and its dominance.

**Fig. 4.**
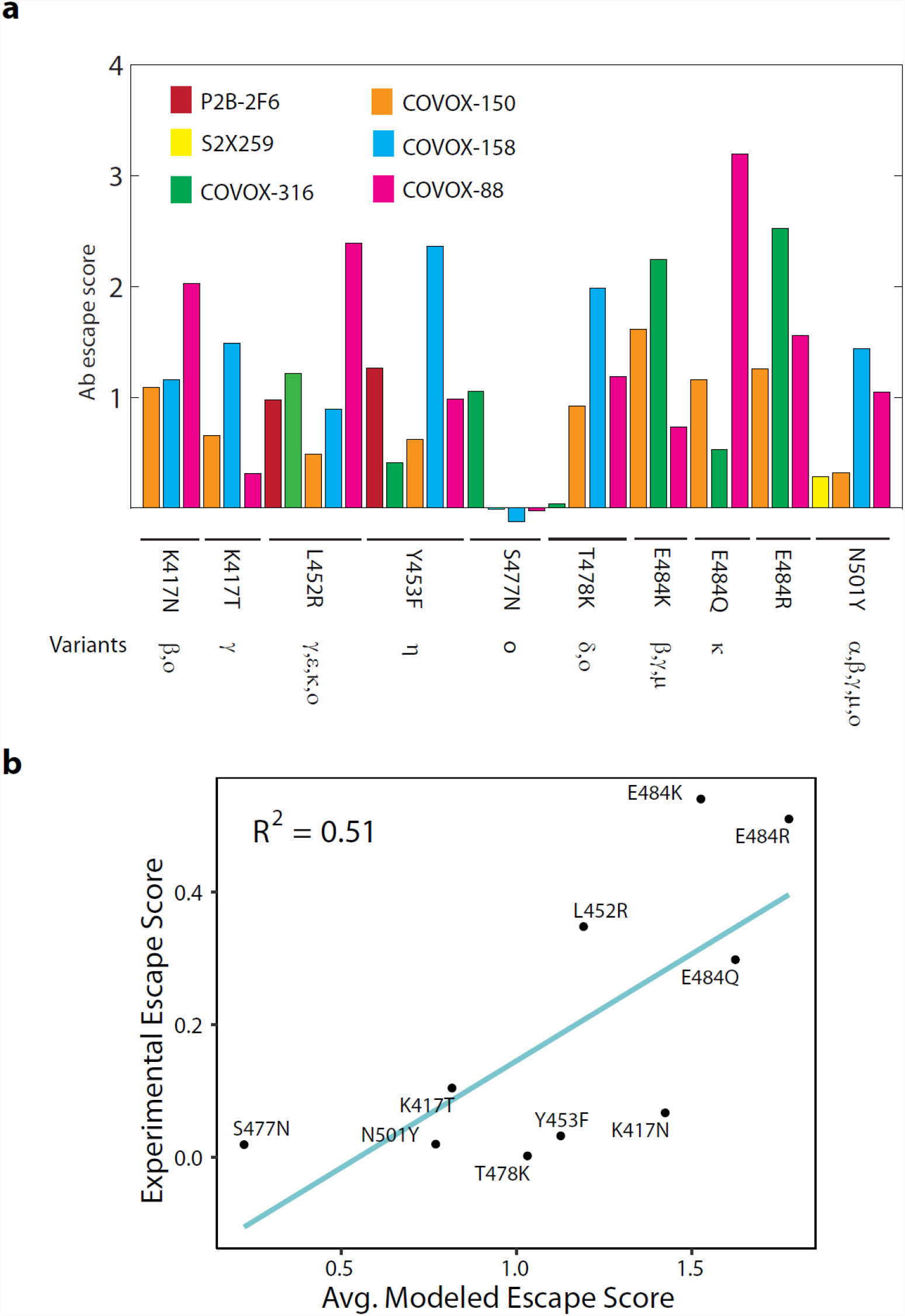
Antibody escape scores of point mutations from non-Omicron variants. **a** Escape scores of point mutations for six neutralizing antibodies targeting S RBD. Occurrence of mutations in different variants is indicated. **b** Correlation between predicted and experimentally measured antibody escape scores. The two approaches used different sets of antibodies targeting S RBD. The experimental score is the sum of contributions from ten antibodies.

A further test of predicted antibody escape scores is to compare with those measured experimentally using deep mutagenesis scanning^17^. The experimental method used ten antibodies targeting S RBD, but the specific antibodies do not overlap with those used in structural modeling. Still, this comparison is meaningful because many S RBD-targeting antibodies bind the crucial S RBM region, as demonstrated by many solved complexes (Fig. 3)^5,14^. By using antibody escape scores for groups of antibodies, we find that the predicted scores are moderately correlated (R^2^=0.51) with those from deep mutagenesis for single mutations (Fig. 4b). As shown, E484K/Q/R and L452R have high antibody escape scores, whereas S477N has a low score. Collectively, our predicted antibody responses to mutations in variants indicate broad qualitative agreement with experimental studies.

### Quantitative antibody escape assessment shows the Omicron S RBD is resistant to most neutralizing antibodies due to unfavorable electrostatic interactions

Although recent experimental works have examined antibody escape by the Omicron variant^4,5^, computational modeling allows identification of specific energetic contributors of antibody resistance. Our antibody escape measure predicted that the Omicron S RBD with 15 mutations can escape all antibodies except S2X259 whose target site is outside of S RBM (Fig. 5a). The antibody escape scores for 5 out of the 6 neutralizing antibodies examined are in the upper range (scores of 2 to 4) of those predicted for single mutations from previous variants (Fig. 4a). In fact, computed affinities show that Omicron Spike cannot form complexes (i.e., ΔG>0) with these tested antibodies. In contrast, the Omicron mutations have a negligible effect on S2X259’s binding to S RBD. These results are in overall agreement with a recent experimental antibody escape study showing that most neutralizing antibodies targeting S RBM are ineffective against Omicron and that effective antibodies target sites outside of S RBM^5^.

**Fig. 5.**
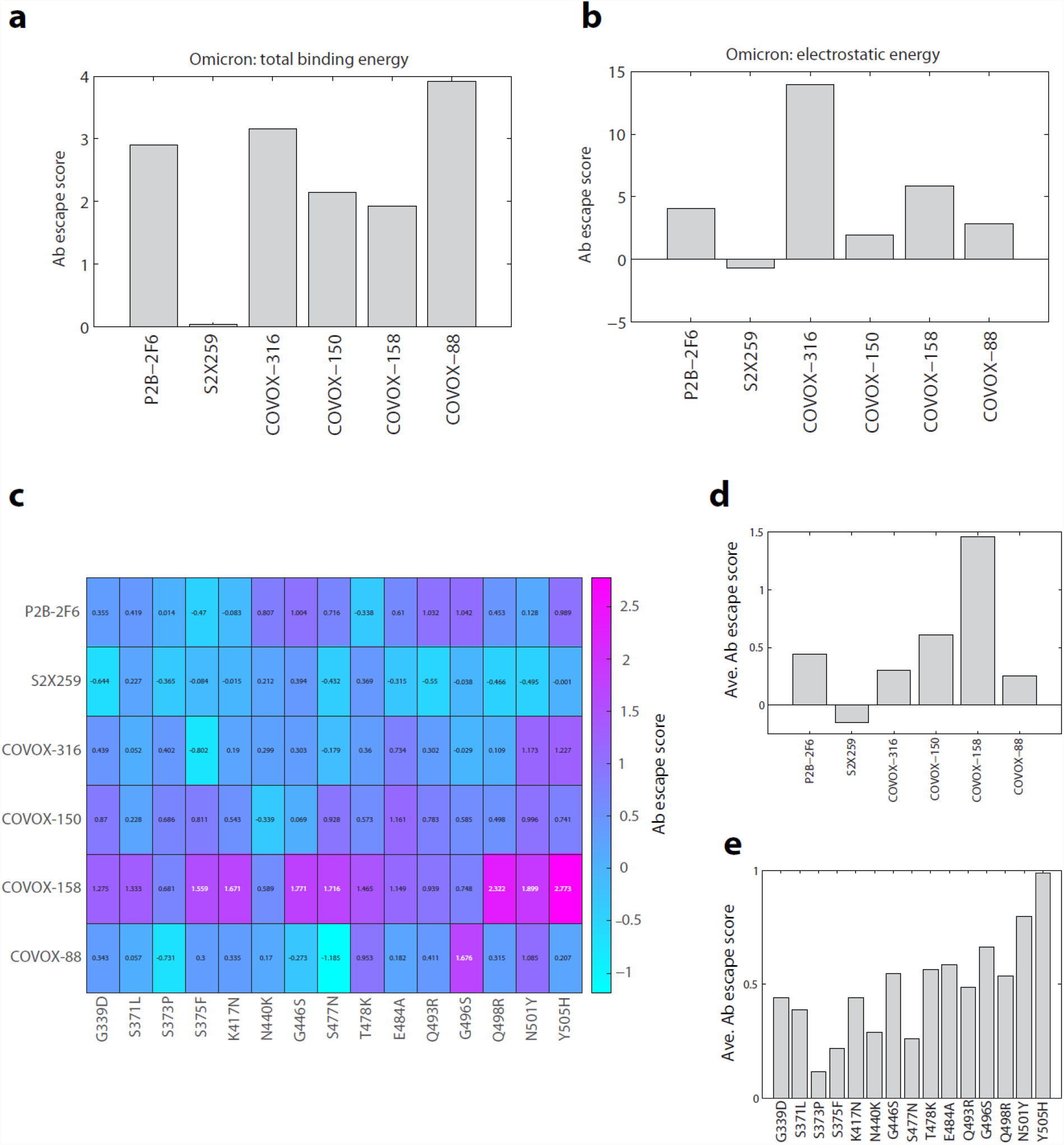
Antibody escape scores of Omicron S RBD mutations. **a** Antibody escape scores of six antibody-Omicron S RBD complexes. **b** Electrostatic component of antibody escape scores of six antibody-Omicron S RBD complexes. **c** Antibody escape score matrix of six antibodies against 15 individual Omicron mutations. **d** Average antibody escape scores of six antibodies induced by individual Omicron mutations (derived from **c**). **e** Antibody escape scores of individual Omicron mutations averaged over six antibodies.

To unravel the energetic factors causing antibody resistance, we examined the major energy components contributing to binding affinity, including short-range van der Waals, electrostatic, and entropy terms^22^. We then computed the antibody escape scores for each energy component (i.e., substituting ΔG_ref_ and ΔG_mut_ with component energies). For the electrostatic energy component, which includes the effects of protein charge and ionic interactions among Spike, antibody and counter-ions, we again observed the same pattern of antibody affinity destabilization by Omicron (Fig. 5b): P2B-2F6 and COVOX-316,-150,-158,-88 have unfavorable electrostatic binding energies, but the effect on S2X259 is nearly unchanged. Electrostatic destabilization is especially strong (score of ∼15) for COVOX-316, which has a positive electrostatic surface like S RBM (Fig. 3b,c). The destabilizing effects of electrostatics are also significant for P2B-2F6 and COVOX-158 (scores of ∼5). Using the same analysis, the effects of other energy components on overall affinity are considerably less. In particular, the changes in escape scores due to van der Waals energy are less than 10%, and there is no specific direction of influence. The entropy factor also contributes to antibody resistance but the effects are weak (escape score of 0.4 for COVOX-88 and much less for other antibodies). This suggests that Omicron mutations introduce some entropic costs to antibody binding. Overall, our quantitative affinity assessment implies that electrostatic interactions are the main contributor to antibody escape by Omicron. This confirms our assessment based on electrostatic surfaces of Spike-antibody complexes (Fig. 3).

To further analyze the contribution of each Omicron mutation to antibody escape, we computationally predicted the escape scores for all combinations of six antibodies and 15 single S RBD mutations (Fig. 5c). As shown, antibody response varies significantly across the 15 mutations and six antibodies. However, a couple of response patterns emerged. Mutation-averaged escape scores show that all antibodies except S2X259 are resisted by individual Omicron mutations (Fig. 5d), consistent with scores modeled with full Omicron S RBD (Fig. 5a). These scores are lower than that for combined Omicron mutations (Fig. 5a), indicating the additive effects of individual mutations. The antibody-averaged scores of individual mutations show that all Omicron mutations can escape antibodies to some extent. This conclusion is in agreement with a recent study of 247 antibodies showing that almost all individual Omicron mutations were shown to escape some antibodies^4^. The predicted stronger escapers (scores > 0.5) are G446S, T478K, E484A, Q493R, G496S, Q498R, N501Y and Y505H and the weaker escapers (scores < 0.5) are G339D, S371L, S373P, S375F, K417N and N440K. Some novel Omicron mutations (G339D, S371L, S373P, S375F, Q493R, G496S, Q498R, and Y505H) are thought to have either a neutral or negative selective advantage^2^. Our modeling indicates that these mutations induce a range of positive antibody escape scores, implying they contribute to the overall fitness of the Omicron variant. Additionally, a recently solved Omicron Spike-ACE2 complex shows that Q493R and Q498R also contribute to affinity by forming two new salt bridges with ACE2^11^, a finding that reinforces the role of electrostatics in Omicron’s interactions.

## Discussion

Long-range electrostatic forces play an important role in biomolecular interactions. Many biomolecular complexes form partly due to the presence of complementary electrostatic potential surfaces^9,10^. Here we show that during the course of the SARS-CoV-2 pandemic variant forms of S RBD have been evolving toward a positively charged surface that complements the ACE2 receptor, ensuring favorable association with the cell surface receptors leading to host cell infection (Fig. 6). At the same time, assessment of electrostatic surfaces of neutralizing antibodies isolated early in the pandemic showed that most of the antibodies have positively charged S RBD-recognition surfaces, implying that recent SARS-CoV-2 variants could escape antibody surveillance via unfavorable electrostatic interactions. Using a computational structural modeling method, we showed this to be the case for the Omicron variant using antibodies targeting different S RBD sites. Thus, a plausible scenario is that SARS-CoV-2 variants have been optimizing their S RBM to simultaneously enhance ACE2 recognition and evade antibodies (Fig. 6). The advantage conferred by the electrostatic property of Omicron S RBD may partially account for the variant’s rapid global transmission.

**Fig. 6.**
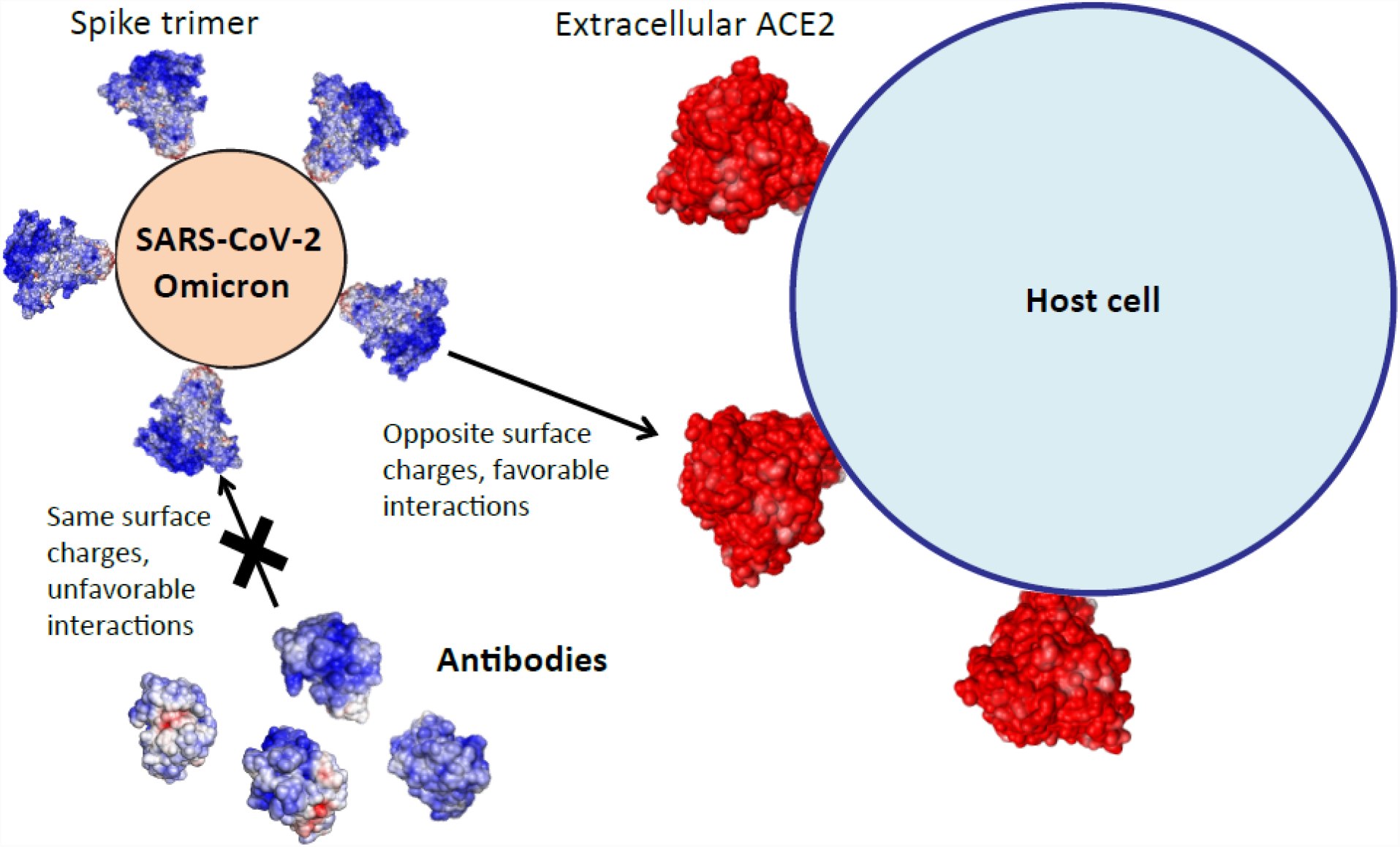
Omicron Spike’s electrostatic potential surface evades most antibodies and promotes ACE2 association for host cell entry.

The evasion of antibodies by the Omicron variant is not complete^4,5^. In particular, antibody S2X259 targeting outside of S RBM (site II) is not destabilized by the Omicron variant, a finding confirmed in a recent screening of antibodies using antibody escape assays^5^. This is due to electrostatic complementarity in the Omicron S RBD-S2X259 complex (Fig. 3b,c). The effectiveness of S2X259 antibody suggests it might be possible to engineer antibodies by exploiting electrostatic complementarity to S RBD target sites, an approach similar to design of ligands capable of binding proteins with high specificity^23^.

The basic character of Omicron S RBD may also impact its interactions with other components of host defense. Specifically, the major components of the mucus system (mucin proteins MUC5AC and MUC5B) shield the respiratory tracts from bacterial and viral infection^24^. Mucin proteins are heavily glycosylated, often with negatively charged terminal sialic acids. Thus the negatively charged polyelectrolytes^25^ in the mucus matrix can in theory provide a more effective shield against Omicron than other variants. This suggests a testable molecular hypothesis for observed attenuated Omicron replication and pathogenicity in respiratory tracts^8^.

The rise of the Omicron family of variants indicates that mutational changes occur in clusters. Analysis of individual mutations may not provide a complete understanding of the biological properties of the new variants^2^. To meet the new challenge, approaches that can probe the collective physical and functional properties are needed. Here we examined how mutational clusters in S RBD of the BA.1 lineage influence ACE2 association and antibody resistance. Thus, future studies on how Omicron mutations collectively affect antibody escape, replication and virulence are needed to provide a broader basis for developing effective intervention strategies.

## Methods

### Prediction of Spike variant structures

The structures were predicted using AlphaFold2, a deep learning approach to protein structure prediction^26^. It enables prediction of Spike structures with mutations, deletions and insertions. The predicted Spike variant structures differed from the solved reference Spike structure (PDB ID: 6vyb) by less than 2 Å.

### Spike-antibody complexes

Six reference Spike-antibody complexes were selected from solved structures based on their structure resolution (<3 Å) and interactions with the 15 Omicron S RBD mutations. They are (PDB ID and Fab name): 7bwj (P2B-2F6), 7m7w (S2X259), 7beh (COVOX-316), 7bei (COVOX-150), 7bek (COVOX-158), 7bel (COVOX-88). The selected antibodies bind multiple sites on S RBD (Fig. 3a). Only the globular Fab domains directly interacting with Spike were retained, i.e., one structured domain each for light and heavy chains.

### Electrostatic potential surfaces of proteins

Electrostatic potential surfaces were computed using APBS v1.5 (at zero ionic concentration)^12^ and visualized using pymol. We used default box dimensions and mesh parameters.

### Structure refinements

Binding affinities of Spike-antibody complexes were calculated based on refined complexes. Structure refinements were performed using Monte Carlo Minimization (MCM) as implemented in Tinker v7.1 molecular modeling package^27^. MCM of each complex was run for six days on four processors on HPC Linux Cluster at NYU Abu Dhabi, with the following parameters: rms gradient < 0.01 kcal/(mol.Å) and move step size of 0.5 Å. We used the all-atom AMBER99/GBSA force field. For affinity calculations, the best energy complexes were used and then subjected to a stringent local minimization (rms gradient of <0.0005 kcal/mol/Å).

### Binding affinities

Affinities of ternary Spike-antibody complexes were computed using molecular modeling methods, as described previously^19,22^. Briefly, the computed binding affinity is a sum of contributions from solvation, van der Waals, electrostatic, and entropic interactions. The electrostatic binding energies were computed using APBS v1.5. All affinities were computed at 37°C and 150 mM of monovalent ions.

## Acknowledgments

This research was carried out on the High Performance Computing resources at New York University Abu Dhabi and New York. We thank Benoit Marchand for his encouragement and support in allocating computational resources for this project. HHG is grateful to Rana Zeine for many useful discussions on immunology.

## Author contributions

H.H.G., K.C.G. and F.P. contributed to plans for the paper. H.H.G. performed structural modeling of electrostatics and antibody escape and wrote the initial draft of the paper. J.Z. predicted structures of Spike variants and performed data analysis. All authors contributed to editing the paper.

## Competing interests

None declared.

## Notes

### Competing Interest Statement

The authors have declared no competing interest.

## References

1. Hadfield, J. et al. Nextstrain: real-time tracking of pathogen evolution. Bioinformatics 34, 4121–4123 (2018).

2. Martin, D.P. et al. Selection analysis identifies unusual clustered mutational changes in Omicron lineage BA.1 that likely impact Spike function. bioRxiv (2022).

3. Gan, H.H., Twaddle, A., Marchand, B. & Gunsalus, K.C. Structural Modeling of the SARS-CoV-2 Spike/Human ACE2 Complex Interface can Identify High-Affinity Variants Associated with Increased Transmissibility. J Mol Biol 433, 167051 (2021).

4. Cao, Y. et al. Omicron escapes the majority of existing SARS-CoV-2 neutralizing antibodies. Nature (2021).

5. Cameroni, E. et al. Broadly neutralizing antibodies overcome SARS-CoV-2 Omicron antigenic shift. Nature (2021).

6. Liu, L. et al. Striking Antibody Evasion Manifested by the Omicron Variant of SARS-CoV-2. Nature (2021).

7. Thomas P. Peacock, J.C.B., Jie Zhou, Nazia Thakur et al. The SARS-CoV-2 variant, Omicron, shows rapid replication in human primary nasal epithelial cultures and efficiently uses the endosomal route of entry. bioRxiv (2022).

8. Shuai, H. et al. Attenuated replication and pathogenicity of SARS-CoV-2 B.1.1.529 Omicron. Nature (2022).

9. Weiner, P.K., Langridge, R., Blaney, J.M., Schaefer, R. & Kollman, P.A. Electrostatic potential molecular surfaces. Proc Natl Acad Sci U S A 79, 3754–8 (1982).

10. McCoy, A.J., Chandana Epa, V. & Colman, P.M. Electrostatic complementarity at protein/protein interfaces. J Mol Biol 268, 570–84 (1997).

11. Mannar, D. et al. SARS-CoV-2 Omicron variant: Antibody evasion and cryo-EM structure of spike protein-ACE2 complex. Science, eabn7760 (2022).

12. Baker, N.A. Poisson-Boltzmann methods for biomolecular electrostatics. Methods Enzymol. 383, 94–118 (2004).

13. Cao, Y. et al. Potent Neutralizing Antibodies against SARS-CoV-2 Identified by High-Throughput Single-Cell Sequencing of Convalescent Patients’ B Cells. Cell 182, 73–84 e16 (2020).

14. Dejnirattisai, W. et al. The antigenic anatomy of SARS-CoV-2 receptor binding domain. Cell 184, 2183–2200 e22 (2021).

15. Corti, D., Purcell, L.A., Snell, G. & Veesler, D. Tackling COVID-19 with neutralizing monoclonal antibodies. Cell 184, 4593–4595 (2021).

16. Weisblum, Y. et al. Escape from neutralizing antibodies by SARS-CoV-2 spike protein variants. Elife 9(2020).

17. Greaney, A.J. et al. Complete Mapping of Mutations to the SARS-CoV-2 Spike Receptor-Binding Domain that Escape Antibody Recognition. Cell Host Microbe 29, 44–57 e9 (2021).

18. Mehra, R. & Kepp, K.P. Structure and Mutations of SARS-CoV-2 Spike Protein: A Focused Overview. ACS Infect Dis 8, 29–58 (2022).

19. Gan, H.H. & Gunsalus, K.C. The Role of Tertiary Structure in MicroRNA Target Recognition. Methods Mol Biol 1970, 43–64 (2019).

20. Flamand, M.N., Gan, H.H., Mayya, V.K., Gunsalus, K.C. & Duchaine, T.F. A non-canonical site reveals the cooperative mechanisms of microRNA-mediated silencing. Nucleic Acids Res 45, 7212–7225 (2017).

21. McCallum, M. et al. Molecular basis of immune evasion by the Delta and Kappa SARS-CoV-2 variants. Science 374, 1621–1626 (2021).

22. Gan, H.H. & Gunsalus, K.C. Tertiary structure-based analysis of microRNA-target interactions. RNA 19, 539–51 (2013).

23. Bauer, M.R. & Mackey, M.D. Electrostatic Complementarity as a Fast and Effective Tool to Optimize Binding and Selectivity of Protein-Ligand Complexes. J Med Chem 62, 3036–3050 (2019).

24. Chatterjee, M., van Putten, J.P.M. & Strijbis, K. Defensive Properties of Mucin Glycoproteins during Respiratory Infections-Relevance for SARS-CoV-2. mBio 11(2020).

25. Sircar, S., Keener, J.P. & Fogelson, A.L. The effect of divalent vs. monovalent ions on the swelling of mucin-like polyelectrolyte gels: governing equations and equilibrium analysis. J Chem Phys 138, 014901 (2013).

26. Jumper, J. et al. Highly accurate protein structure prediction with AlphaFold. Nature 596, 583–589 (2021).

27. Rackers, J.A. et al. Tinker 8: Software Tools for Molecular Design. J Chem Theory Comput 14, 5273–5289 (2018).

